# AHCY inhibition reshapes DNA methylation and transcriptional regulation in colorectal cancer cells

**DOI:** 10.64898/2026.11.26.769437

**Authors:** Iiro Tikkakoski, Konsta Karttunen, Divyesh Patel, Subhamoy Datta, Biswajyoti Sahu, Päivi Pihlajamaa

## Abstract

DNA methylation is an important epigenetic mechanism regulating gene expression, and its maintenance depends on cellular methyl group metabolism. Adenosylhomocysteinase (AHCY) is a key enzyme in the methionine cycle that hydrolyzes S-adenosylhomocysteine (SAH), thereby preventing its accumulation and inhibition of methyltransferase activity. Although AHCY has been implicated in cancer cell proliferation and transcriptional regulation, the effects of its inhibition on genome-wide DNA methylation and their relationship to gene expression remain poorly understood in colorectal cancer (CRC). Here, we investigated the consequences of pharmacological AHCY inhibition in GP5d colorectal cancer cells by integrating metabolomic profiling, nanopore sequencing-based DNA methylation analysis and RNA sequencing. AHCY inhibition resulted in SAH accumulation, widespread DNA hypomethylation and reduced cell proliferation, accompanied by changes in purine metabolism and gene expression. Despite the overall loss of DNA methylation, localized hypermethylation was also observed, particularly at CpG-rich regulatory regions. Transcriptomic analysis revealed changes in proliferation-associated and cell cycle checkpoint pathways, including strong upregulation of the cell cycle regulator *CDKN1A*. Importantly, promoter hypomethylation was significantly enriched among upregulated genes, including genes involved in cell cycle regulation, whereas gene body hypomethylation showed no comparable association with differential gene expression. Together, these findings demonstrate that AHCY inhibition induces extensive metabolic, epigenetic and transcriptional changes in CRC cells and suggest that promoter hypomethylation may contribute to the transcriptional responses associated with reduced cell proliferation.

## Introduction

Methylation reactions are involved in numerous cellular processes, including post-translational modifications and the regulation of gene expression. These reactions depend on the methionine cycle, which generates S-adenosylmethionine (SAM), the principal cellular methyl donor. Following methyl group transfer, SAM is converted to S-adenosylhomocysteine (SAH), which inhibits SAM- dependent methyltransferases. Adenosylhomocysteine hydrolase (AHCY) is the only known enzyme in humans that catalyzes the hydrolysis of SAH into homocysteine and adenosine, thereby preventing SAH accumulation and maintaining cellular methylation capacity (1–3). Thus, AHCY acts as a metabolic gatekeeper, regulating SAH, adenosine and homocysteine levels, and influencing cellular processes such as redox homeostasis and stress response (4). AHCY has been reported to be upregulated in many cancer cell lines (2,5–7), and previous studies have demonstrated that its inhibition affects gene expression in multiple cell lines (2,8–10). However, the epigenetic mechanisms underlying these transcriptional changes, particularly in colorectal cancer (CRC), remain poorly understood. Here, we investigate how AHCY inhibition affects DNA methylation, gene expression, and metabolism in CRC cells.

By regulating SAH accumulation and thereby the activity of SAM-dependent methyltransferases, AHCY can influence methylation of DNA, RNA, and histone proteins, with broad consequences for epigenetic regulation and gene expression. AHCY has been shown to maintain histone modifications associated with both transcriptional activation and repression, such as H3K4me3 and H3K27me3, respectively (11). Cancer cells have also been shown to exploit methionine metabolism through methionine overconsumption, leading to loss of H3K79me2 and impaired CD8^+^ T cell immunity (12). AHCY is also important for RNA methylation (13). For example, N6-methyladenosine (m^6^A) modification of mRNA can influence translation efficiency, mRNA stability and nuclear export (14). Furthermore, AHCY has been shown to form adenosine-dependent dimers that enhance m^6^A methylation and promote lipogenesis and tumorigenesis, whereas mutations that disrupt AHCY dimerization or loss of AHCY suppresses these processes (15).

In addition to histone and RNA methylation, AHCY is required for DNA methylation (1). DNA methylation predominantly occurs at cytosines in CpG dinucleotides and is generally associated with transcriptional repression when present at gene promoters. Although promoters frequently overlap CpG islands, these regions typically remain unmethylated, partly through the binding of transcription factors that restrict access of DNA methyltransferases (DNMT). The distribution of CpG sites and transcription factor binding sites varies between genes and transcript isoforms, potentially influencing their sensitivity to changes in DNA methylation (16,17). Previous studies have shown that combined inhibition of DNMTs and histone deacetylases (HDAC) induces DNA hypomethylation and transcriptional activation of transposable elements (18). Furthermore, AHCY is overexpressed in several cancer cell lines, suggesting a potential role in regulating transcriptional activity in cancer (6). Importantly, metabolites such as SAM, flavin adenine dinucleotide (FAD), and nicotinamide adenine dinucleotide (NAD) can directly modulate the activity of epigenetic enzymes by acting as cofactors or substrates (19). Thus, alterations in AHCY activity may affect gene regulation through multiple interconnected metabolic and epigenetic mechanisms.

Aberrant DNA methylation patterns are a common feature of CRC progression. A subset of CRCs exhibit a distinct methylation pattern known as the CpG island methylator phenotype (CIMP), characterized by widespread hypermethylation of promoter-like regions and CpG islands (20,21). CIMP has been associated with BRAF mutations, mismatch repair deficiency, and increased metabolic activity, including the one-carbon metabolism and the methionine cycle (22). Together, these observations suggest an important role for altered DNA methylation in CRC progression (20–22). Emerging evidence also suggests that epigenetic alterations, including changes in DNA methylation, changes may contribute to immune evasion and treatment resistance in CRC (23).

CRC is one of the leading causes of cancer-related deaths worldwide (24). One-carbon metabolism has long been targeted in CRC treatment, notably by the widely used chemotherapeutic agent 5-fluorouracil (5-FU), which inhibits thymidylate synthase and thereby disrupts nucleotide synthesis (25). More recently, AHCY has emerged as a potential therapeutic target, with pharmacological inhibition shown to reduce tumor burden in a mouse model (6). However, AHCY also plays important roles in normal cellular functions, as demonstrated by the early embryonic lethality observed in AHCY knockout mice (26,27). Furthermore, AHCY is highly expressed in pluripotent embryonic stem cells and has been implicated in maintaining their transcriptional activity (28). Together, these findings highlight the importance of AHCY in normal development and suggest that its therapeutic inhibition may require careful consideration of potential effects on normal tissues.

Previous studies have demonstrated that AHCY regulates gene expression and proliferation in cancer cells, although its requirement for cell survival varies between cellular contexts (2,8,29). However, the epigenetic mechanisms underlying these transcriptional changes, particularly the contributions of DNA methylation, remain poorly characterized in CRC. Nanopore sequencing has emerged as a powerful long-read sequencing technique that enables the direct detection of DNA methylation from native DNA (30). Here, we integrate nanopore-based DNA methylation profiling with RNA sequencing and metabolomics to investigate the effects of AHCY inhibition in GP5d CRC cells. We demonstrate that AHCY inhibition induces widespread DNA hypomethylation, alters purine metabolism, and reduces cell proliferation. Furthermore, we identify an association between promoter hypomethylation and the upregulation of genes involved in cell-cycle regulation, providing insight into the metabolic and epigenetic consequences of AHCY inhibition in CRC.

## Results

### AHCY inhibition reduces cell proliferation and alters purine metabolism

To investigate the cellular consequences of AHCY inhibition in colorectal cancer, we treated GP5d cells with DZNep, a pharmacological AHCY inhibitor (31). AHCY inhibition leads to SAH accumulation, reduced cellular methylation capacity, and impaired adenosine production through the methionine cycle (Fig. 1A). We then performed metabolite profiling, RNA sequencing (RNA-seq) and nanopore sequencing (Fig. 1B) to characterize the metabolic, transcriptional, and DNA methylation changes following DZNep treatment (Fig. 1B). DZNep treatment significantly reduced GP5d cell proliferation compared with vehicle-treated controls (Fig. 1C). This finding is consistent with previous CRISPR screening data from DepMap showing that AHCY depletion impairs cell proliferation (29,32), as well as a previous study demonstrating reduced growth of mouse CRC organoids following AHCY inhibition (6).

**Fig. 1.**
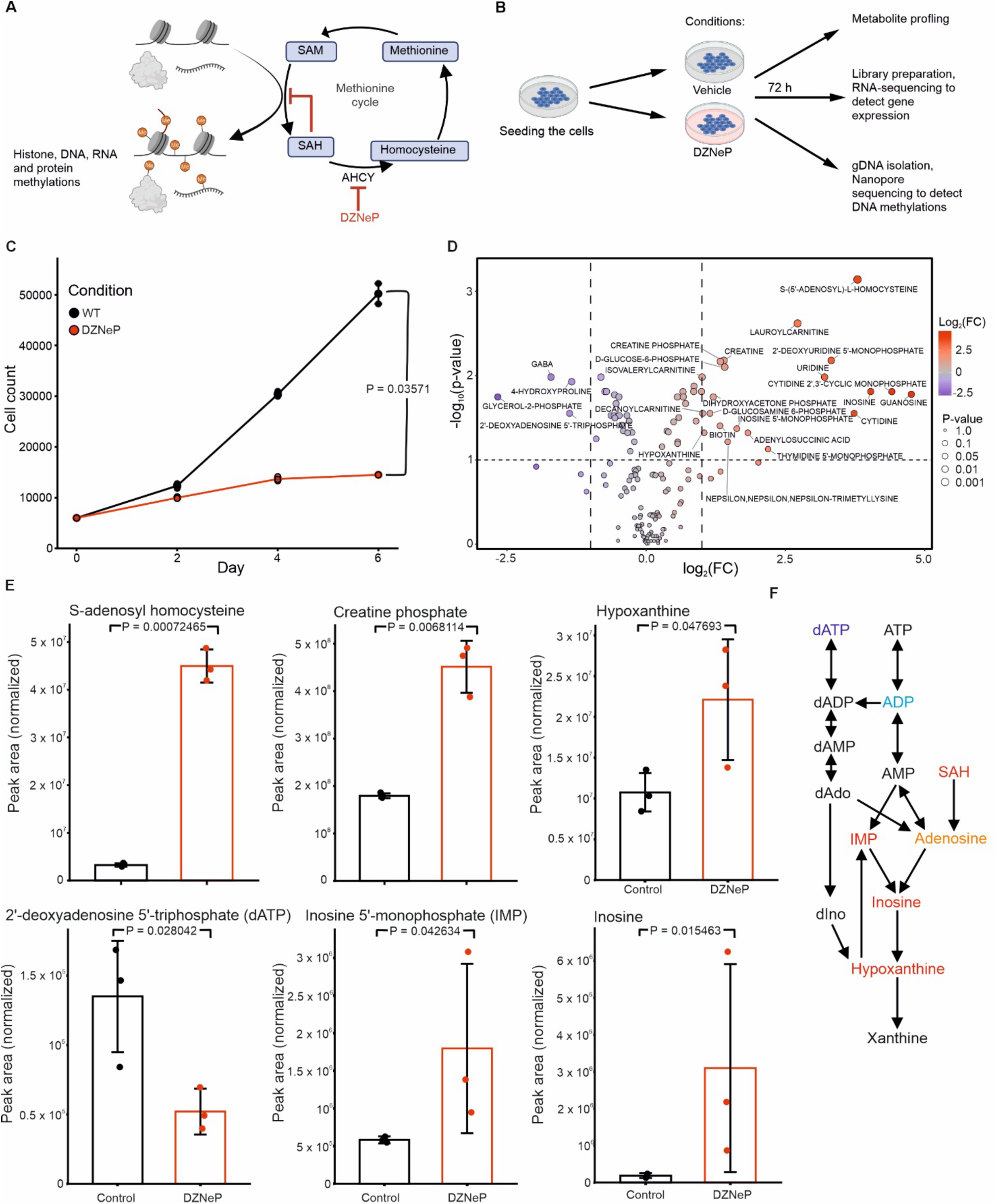
AHCY inhibition reduces cell proliferation and alters purine metabolism. (**A**) Schematic of the methionine cycle. Methionine is converted to SAM, which serves as a methyl donor for DNA, histones or protein modifications. Following the methyl group transfer, SAM is converted to SAH, which inhibits SAM-dependent methylation reactions. AHCY catalyzes the hydrolysis of SAH to homocysteine and adenosine. Homocysteine can subsequently be converted back to methionine. DZNep directly inhibits AHCY. (**B**) Experimental workflow. GP5d cells were treated with DZNep or vehicle for 72 hours, followed by metabolite profiling, RNA-seq and Nanopore sequencing. (**C**) GP5d cell numbers in vehicle- and DZNep-treated conditions measured at two-day intervals. Dots represent technical replicates (n = 3). Cell numbers differed significantly between conditions on day 6 (Welch two sample t-test, P = 0.03571) (**D**) Volcano plot of metabolite profiling results. The x-axis represents the log_2_ fold change (log_2_FC) of metabolite abundance in DZNep-treated cells compared to control, and y-axis represents the –log_10_ p-value. Four metabolites were significantly decreased and 15 metabolites were significantly increased in DZNep condition. (**E**) Bar plots showing selected significantly altered metabolites. The y-axis shows normalized metabolite peak area. Coloured dots represent individual replicates, and error bars represent ±1 standard deviation. P, adjusted p-value (t-test). (**F**) Simplified schematic of purine metabolism showing pathway from ATP and dATP to xanthine and the production of adenosine from SAH in the methionine cycle. dAdo, deoxyadenosine; dIno, deoxyinosine. Bidirectional arrows indicate reversible reactions catalysed by distinct enzymes. Metabolite names are coloured according to the observed direction of change and statistical significance: dark blue, decreased (p < 0.05); light blue, decreased (p < 0.1); orange, increased (p < 0.1); red, increased (p < 0.05).

Metabolite profiling following AHCY inhibition revealed several significantly altered metabolites. Many were associated with the methionine cycle and purine and pyrimidine metabolism, while individual metabolites involved in energy storage (creatine and phosphocreatine), and lipid and glucose metabolism were also increased (Fig. 1D). As expected, SAH levels increased following DZNep treatment, consistent with inhibition of AHCY activity. Notably, several purine metabolites, such as inosine and inosine monophosphate (IMP), were increased, whereas deoxyadenosine triphosphate (dATP) was significantly decreased (Fig. 1E).

Further examination of purine metabolism revealed a broader pattern of changes in adenine nucleotide metabolism. ATP, ADP, AMP, and dATP levels were decreased, whereas adenosine and its downstream metabolites, including inosine, IMP, and hypoxanthine, were increased (Fig. 1E, Fig. 1F). These changes were most pronounced for dATP, inosine, IMP, and hypoxanthine, which showed statistically significant differences. The metabolic pathways connecting adenine nucleotides to adenosine and downstream purine degradation products are well established (33–35). Together, our findings suggest that AHCY inhibition induces a shift from methionine-cycle-derived adenosine production toward compensatory adenosine production through alternative sources, such as ATP and dATP degradation. Since dATP is essential for DNA synthesis, its reduced levels may contribute to the impaired proliferation observed following DZNep treatment (36). Conversely, reduced cell proliferation may itself lead to lower dATP levels through decreased nucleotide demand and synthesis.

### AHCY inhibition induces widespread DNA hypomethylation

To investigate the effects of AHCY inhibition on DNA methylation, we performed nanopore sequencing of vehicle- and DZNep-treated GP5d cells. Nanopore sequencing enables direct detection of DNA methylation from native DNA, allowing genome-wide analysis of CpG methylation and identification of differentially methylated regions (DMR) (30,37).

We first examined the distribution of CpG methylation at the genome-wide level. In control cells, CpG methylation displayed a bimodal distribution, with most CpGs either highly methylated or unmethylated with an average methylation level of 64.9% (Fig. 2A). Following DZNep treatment, the distribution shifted toward lower methylation levels, with a reduced fraction of highly methylated CpGs and an increased fraction of CpGs with intermediate methylation levels. Consequently, the average CpG methylation level decreased to 49.9% (Fig. 2A). We then identified differentially methylated regions (DMR), defined as regions of at least 100 bp containing at least four CpGs and a two-sided Fisher’s test FDR below 0.05 between the conditions. In total, 321,427 DMRs were detected, of which 318,986 were hypomethylated and only 2,441 were hypermethylated (Fig. 2B,C). The minority of DMRs that gained methylation were strongly enriched in partially methylated domains (PMD): 77.0% of hypermethylated DMRs overlapped PMDs, compared with 43.0% of hypomethylated DMRs (Fisher’s exact test, OR = 4.42, 95% CI 4.02–4.87, P = 1.3 × 10⁻²⁵³). PMDs are late-replicating regions that are particularly prone to methylation loss over successive cell divisions (38), consistent with the substantial overlap of hypomethylated DMRs with these domains. The preferential overlap of hypermethylated DMRs with PMDs, on the other hand, is consistent with the pattern described in CRC cells upon DNMT1 loss, where focal methylation gains amid global hypomethylation arise through region-specific de novo methyltransferase activity (39).

**Fig. 2.**
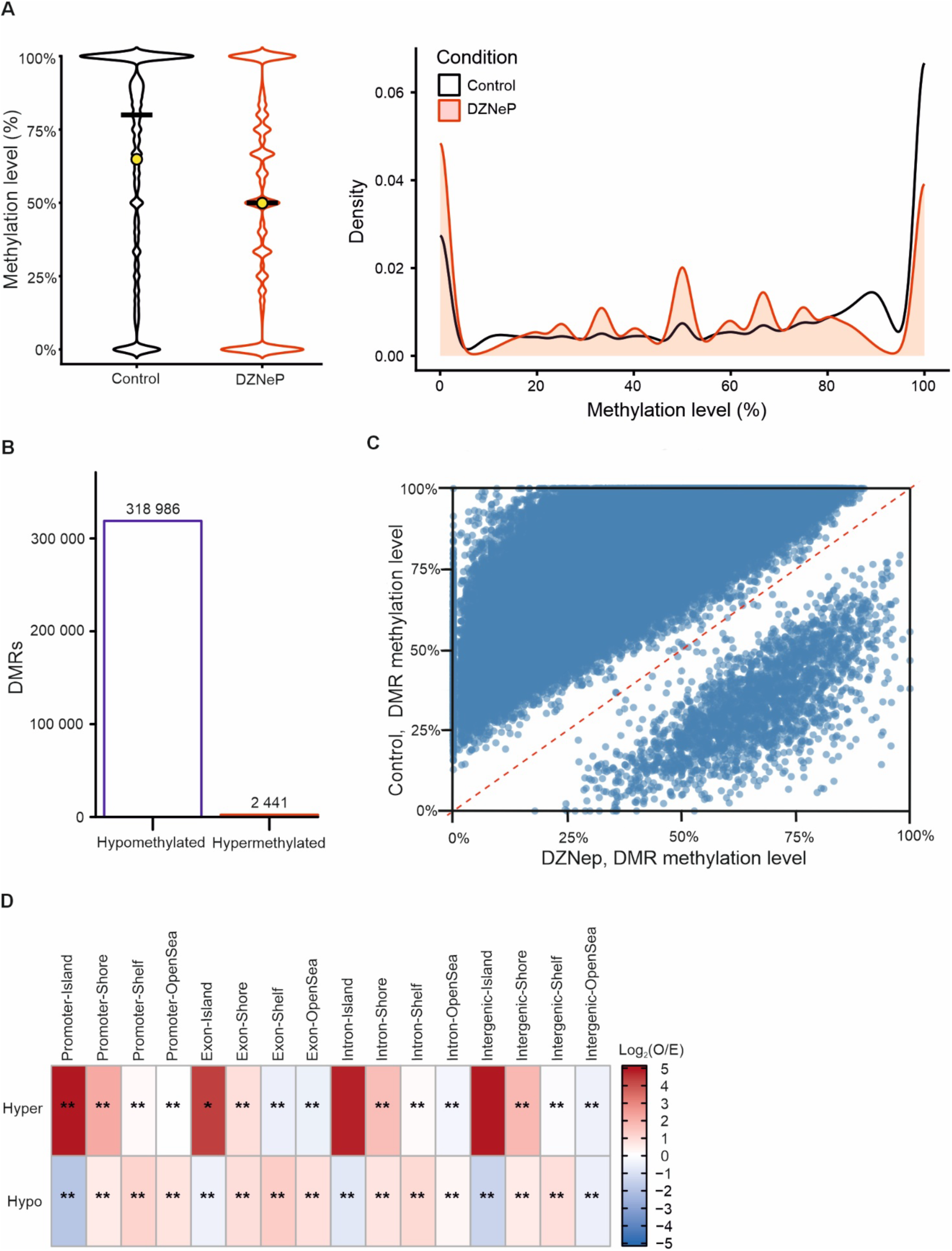
AHCY inhibition induces widespread DNA hypomethylation. (**A**) Distribution of CpG methylation levels in vehicle- and DZNep-treated GP5d cells, shown as violin plots (left) and density plots (right). The black bar indicates the median and the yellow circle indicates the mean methylation level. Mean CpG methylation decreased from 64.9% in control cells to 49.9% following DZNep treatment. (**B**) Numbers of hypo- and hypermethylated DMRs following DZNep treatment. (**C**) Scatter plot showing the average methylation level of each DMR in control (y-axis) and DZNep-treated (x-axis) cells. The red diagonal line indicates equal methylation between conditions. Points above the line represent hypomethylated DMRs following DZNep treatment, whereas points below the line represent hypermethylated DMRs. **(D)** Observed vs expected enrichment of hyper- and hypomethylated DMRs across genomic elements (promoters, exons, introns and intergenic regions) stratified by CpG island proximity (islands, shores, shelves, and open sea). Overrepresentation, red; underrepresentation, blue. Asterisks indicate statistical significance (p-values were obtained from Monte Carlo simulations. *, p-value < 0.05; **, p-value < 0.01). Promoters were defined as regions 2500 bp upstream and 500 bp downstream of TSS. CpG shores and shelves were defined as regions within 2000 bp and regions 2,000–4,000 bp from CpG islands, respectively.

We next quantified DMR enrichment across annotated genomic regions by comparing the observed number of DMRs in each category to the number expected by chance. Hypermethylated DMRs were strongly and significantly enriched at CpG islands of promoters and exons (Fig. 2D). Enrichment at intronic and intergenic CpG islands was of similar magnitude (Fig. 2D), but did not reach statistical significance, likely owing to the small number of DMRs in these categories. Conversely, hypomethylated DMRs were underrepresented at CpG islands where methylation levels are generally low, but showed enrichment at CpG shores and shelves (Fig. 2D).

Together, these findings demonstrate that AHCY inhibition induces extensive genome-wide DNA hypomethylation, accompanied by localized methylation gains that preferentially occur in partially methylated domains and at CpG islands.

### AHCY inhibition alters proliferation-associated transcriptional programs

To investigate the transcriptional changes associated with AHCY inhibition, we performed RNA-seq of vehicle- and DZNep-treated GP5d cells. We identified 1,965 differentially expressed genes (DEG), of which 770 were upregulated and 1,195 downregulated following DZNep treatment (Fig. 3A). Among the most significantly altered genes, ID2 showed the most significant upregulation, whereas *TXNIP* was the most significantly downregulated gene (Fig. 3A). ID2 has been implicated in cell cycle regulation through its interaction with transcriptional regulators and the RB pathway (40). *TXNIP* expression is reduced in multiple cancer cell lines and it has roles in redox homeostasis, cellular metabolism and has been described as a tumor suppressor gene (41). *AHCY* expression itself was not altered following DZNep treatment (Fig. 3A).

**Fig. 3.**
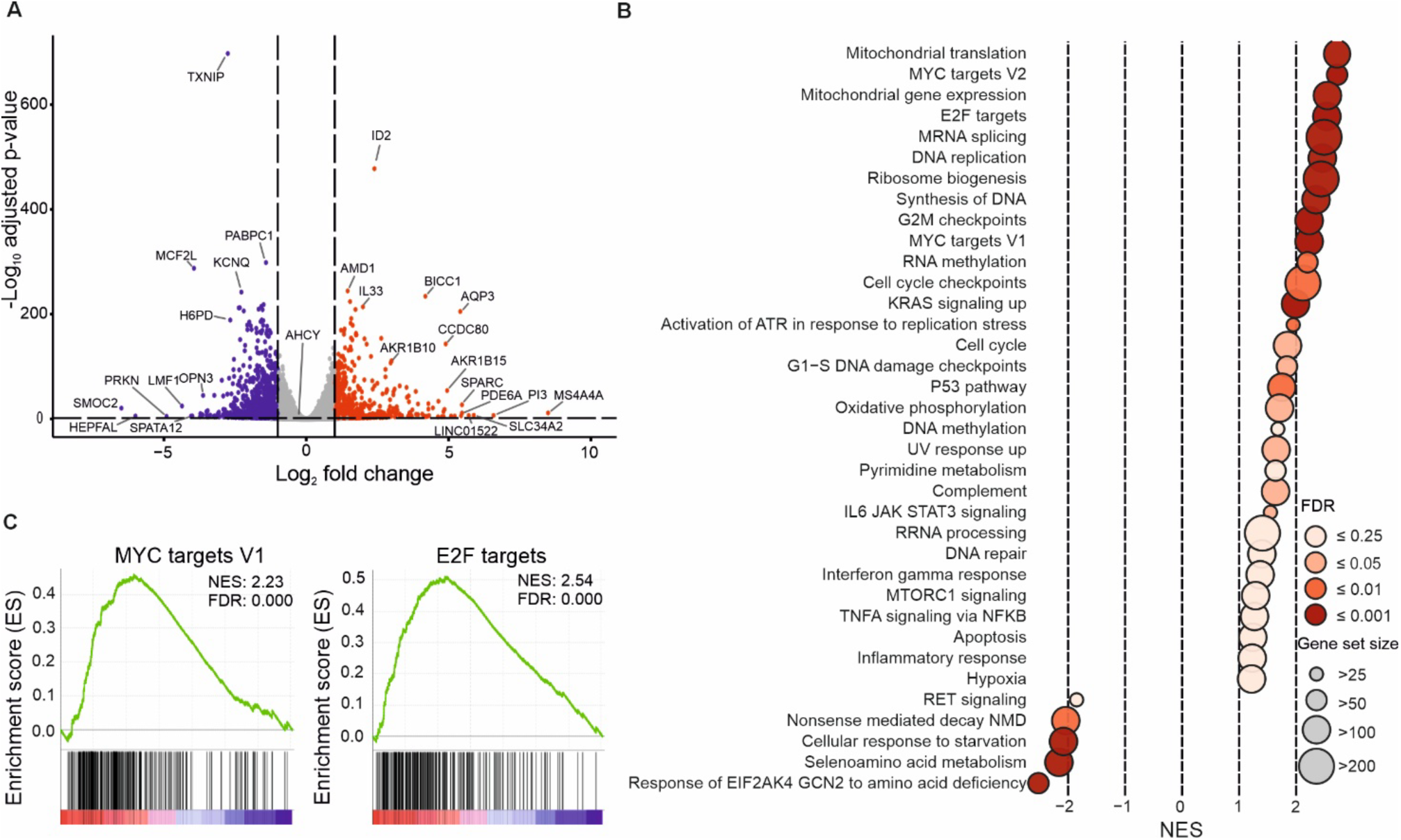
AHCY inhibition alters proliferation-associated transcriptional programs. (**A**) Volcano plot from RNA-seq data showing differential gene expression in DZNep-treated GP5d cells compared to vehicle control. The x-axis represents log_2_ fold change of genes and y-axis -log_10_ adjusted p-values. Significantly upregulated genes are shown in red and downregulated in blue (log_2_ fold change > 1, adjusted p-value < 0.05), with 770 genes up- and 1195 downregulated genes followed by DZNep treatment. (**B**) Gene set enrichment analysis (GSEA) of RNA-seq data comparing DZNep-treated and control cells. Each bubble represents a gene set, with its normalized enrichment score (NES) shown on the x-axis. Darker red indicates greater statistical significance (lower FDR). (**C**) Enrichment plots for selected gene sets. Genes in red areas are upregulated and genes in blue areas are downregulated in DZNep condition.

To identify the biological processes affected by AHCY inhibition, we performed gene set enrichment analysis (GSEA). Interestingly, despite the reduced proliferation observed following DZNep treatment, gene sets associated with cell proliferation, such as MYC targets, E2F targets, DNA replication and pyrimidine metabolism, were significantly enriched among genes upregulated in DZNep-treated cells (Fig. 3B,C). At the same time, gene sets associated with cell cycle checkpoints, the G1/S DNA damage checkpoint and the p53 pathway were also enriched (Fig. 3B), consistent with previous findings (8). Together, this suggest that AHCY inhibition induces a complex transcriptional response involving both proliferation-associated programs and cell cycle checkpoint pathways, despite an overall reduction in cell proliferation.

In addition to proliferation-related pathways, gene sets related to mitochondrial function, oxidative phosphorylation and ribosome biogenesis were enriched following DZNep treatment (Fig. 3B), indicating broader changes in metabolic and biosynthetic transcriptional programs (42–44). These changes are consistent with the alterations observed in adenine nucleotide metabolism observed in our metabolomic analysis (45,46). Changes in genes associated with creatine metabolism were also observed, consistent with the increased phosphocreatine levels detected by metabolomic profiling (c.f. Fig. 1E). However, the expression data did not provide clear evidence of increased creatine or phosphocreatine synthesis. In addition, a gene set associated with ATR activation in response to replication stress was enriched following DZNep treatment, which is consistent with the nucleotide pool stress observed in metabolomic profiling (47,48).

Examination of individual genes within the enriched gene sets revealed that *CDKN1A*, encoding the cyclin-dependent kinase inhibitor p21, was among the most strongly upregulated genes (see Fig. 4B). Given the key role of *CDKN1A* in regulating cell cycle progression at the G1/S and G2/M transitions (46) (49), its upregulation may contribute to the reduced proliferation observed following DZNep treatment. Together, these findings indicate that AHCY inhibition induces widespread transcriptional changes involving both proliferation-associated and cell cycle checkpoint pathways, with *CDKN1A* upregulation potentially contributing to the observed reduction in cell proliferation.

**Fig. 4.**
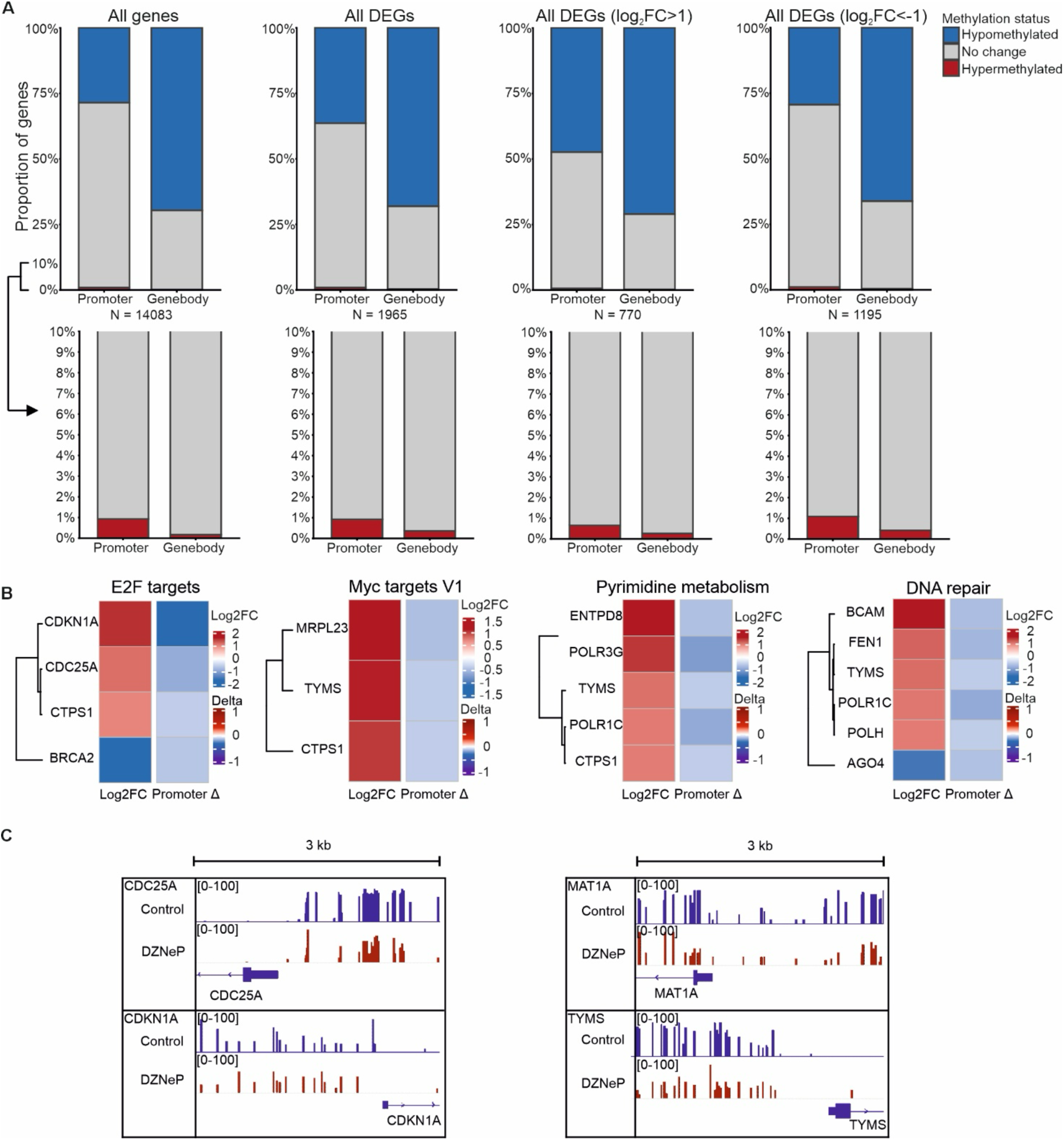
**Promoter hypomethylation is enriched among upregulated genes upon AHCY inhibition**. (**A**) Proportions of genes showing promoter or gene body hypo- and hypermethylation following DZNep treatment, shown for all analyzed genes, all DEGs, upregulated DEGs and downregulated DEGs. The number of genes in each category is indicated by n. The lower panels show an enlarged view of the bottom 10% of the corresponding plots to visualize the smaller proportions of hypermethylated genes. Promoter hypomethylation was significantly enriched among upregulated genes (OR 2.38, 95% CI 2.05–2.76, Fisher’s exact test, P = 5.424*10^-30^), but not among downregulated genes (OR 1.04, 95% CI 0.91–1.19, Fisher’s exact test, P = 0.5248). Promoter hypermethylation was not enriched among upregulated genes (OR 0.68, 95% CI 0.22–1.65, P = 0.5604) nor downregulated genes (OR 1.19, 95% CI 0.61–2.12, P = 0.5278). (**B**) Gene expression and promoter methylation changes in selected gene sets. Left column shows RNA-seq log_2_ fold changes and right column shows changes in DNA methylation (delta value). Only genes with significant expression changes (absolute log2 fold change > 1, adjusted P < 0.05) and promoter methylation changes (absolute methylation difference > 0.1, FDR < 0.05) are shown. (**C**) Genome browser views showing CpG methylation at selected genes following DZNep treatment. The upper and middle tracks show the proportions of methylated CpG reads in control and DZNep-treated cells, respectively, and the lower track shows gene annotations. Promoter was defined as a region from 500 bp downstream and 2500 bp upstream from TSS.

### Promoter hypomethylation is enriched among upregulated genes upon AHCY inhibition

To investigate the relationship between DNA methylation and gene expression changes following AHCY inhibition, we integrated the nanopore methylation data with our RNA-seq results. We first examined methylation changes at promoters and gene bodies independently of DMRs by calculating differences in CpG methylation levels between conditions. Across all analyzed genes, approximately one-quarter of promoters and more than 60% of gene bodies showed reduced methylation following DZNep treatment (Fig. 4A). However, when focusing on DEGs, we observed that the proportion of hypomethylated promoters was nearly twice as high among upregulated genes compared to all analyzed genes (Fig. 4A). This enrichment was statistically significant (OR = 2.38, 95% CI 2.05– 2.76, Fisher’s exact test, P = 5.424 × 10⁻³⁰), whereas no significant enrichment was observed among downregulated genes, or for promoter hypermethylation among either upregulated or downregulated genes (Fig. 4A). In contrast, the proportion of genes showing gene body hypomethylation was similar across expression categories (Fig. 4A). These results indicate that promoter hypomethylation is preferentially associated with gene upregulation following AHCY inhibition, whereas promoter hypermethylation shows no significant association with differential gene expression.

We next examined the relationship between promoter methylation and gene expression changes within the gene sets identified by GSEA. Focusing on genes with significant changes in both promoter methylation and expression, we found that majority showed promoter hypomethylation accompanied by increased expression (Fig. 4B). This pattern was observed across several positively enriched gene sets, including those associated with cell proliferation and nucleotide metabolism. Examination of individual loci using IGV revealed reduced methylation near the transcription start sites (TSS) of *CDKN1A* and *MAT1A*, with similar changes observed farther from the TSSs of *TYMS* and *CDC25A* (Fig. 4C). Together, these findings suggest that promoter hypomethylation may contribute to the transcriptional changes induced by AHCY inhibition, particularly in genes involved in cell cycle regulation and nucleotide metabolism.

## Discussion

Here, we investigated the effects of AHCY inhibition on cellular metabolism, DNA methylation and gene expression in GP5d colorectal cancer cells. DZNep treatment resulted in SAH accumulation, widespread DNA hypomethylation and reduced cell proliferation, accompanied by substantial changes in nucleotide metabolism and transcriptional regulation. Importantly, integration of nanopore methylation and RNA-seq data revealed that promoter hypomethylation was significantly enriched among upregulated genes, suggesting that altered DNA methylation may contribute to the transcriptional responses to AHCY inhibition.

A recent study of cancer cachexia demonstrated that disruption of methionine cycle metabolism can induce promoter hypomethylation and upregulation of Ddit4/REDD1 in skeletal muscle (50), supporting a connection between methionine cycle metabolism, DNA methylation and transcriptional regulation. Interestingly, although AHCY inhibition induced widespread DNA hypomethylation in our study, hypomethylated DMRs were underrepresented at promoter CpG islands, whereas hypermethylated DMRs were enriched at these regions. This suggests that different genomic regions vary in their susceptibility to methylation changes following AHCY inhibition. Many DMRs were also located outside promoters, where their consequences for gene regulation remain less clear. In addition, changes in purine metabolites and enrichment of gene sets associated with mitochondrial energy metabolism suggest that AHCY inhibition affects broader metabolic processes. Given the roles of adenosine and adenine nucleotides in cellular energy metabolism and nucleotide synthesis (8,35,36), these changes may contribute to the cellular response to AHCY inhibition.

Our RNA-seq results were partly consistent with previous studies investigating AHCY depletion in different cancer cell lines (2,8). In SW480 colorectal cancer cells, shRNA-mediated AHCY silencing resulted in MYC upregulation and altered expression of genes involved in Wnt and STAT3 signaling, cell cycle regulation, cytoskeletal organization and cellular migration (2). In HepG2 liver cancer cells, AHCY knockdown was associated with reduced proliferation, cell cycle arrest, increased DNA damage responses and altered expression of genes involved in cell cycle checkpoints, DNA repair and stress signaling (8). These studies suggest that the transcriptional consequences of AHCY depletion vary between cellular contexts. In our GP5d cells, the enrichment of MYC targets and other proliferation-associated gene sets resembled some of the findings in SW480 cells, whereas the reduced proliferation and enrichment of checkpoint-associated pathways were consistent with observations in HepG2 cells.

Interestingly, despite the enrichment of proliferation-associated gene sets, DZNep treatment reduced GP5d cell proliferation. This apparent discrepancy may reflect the simultaneous upregulation of cell cycle checkpoint pathways that limit cell proliferation. Previous studies have shown that treatment with 1 µM DZNep induces accumulation of OCI-AML3 and HL-60 cells in the G0/G1 phases of the cell cycle (51). A similar response may occur in GP5d cells, as *CDKN1A*, encoding the cell cycle inhibitor p21, was strongly upregulated following DZNep treatment. We also observed reduced DNA methylation near the *CDKN1A* TSS, consistent with previous findings linking *CDKN1A* promoter demethylation to increased expression (52). Furthermore, DZNep has been reported to induce a p53/p21-mediated senescence response associated with DNA damage signaling (53). Together, these observations suggest that *CDKN1A* upregulation may contribute to the reduced proliferation following AHCY inhibition, although direct cell cycle measurements would be needed to determine whether DZNep induces cell cycle arrest in GP5d cells.

The combined metabolomic and transcriptomic results suggest that changes in nucleotide metabolism may also contribute to the cellular response to AHCY inhibition. DZNep treatment altered several purine metabolites and significantly reduced dATP levels, while ATP, ADP and AMP showed nonsignificant decreases. These changes could potentially affect DNA replication and cellular energy metabolism, consistent with the enrichment of gene sets associated with DNA replication, cell cycle checkpoints and mitochondrial function. However, the observed changes in nucleotide levels may also reflect altered metabolic demands associated with reduced cell proliferation. Together with the methylation-associated transcriptional changes, these findings suggest that both metabolic and epigenetic consequences of AHCY inhibition may contribute to the reduced proliferation of GP5d cells.

An important consideration is that the transcriptional effects of AHCY inhibition are unlikely to be explained by DNA methylation changes alone. Although promoter hypomethylation was enriched among upregulated genes, some genes showed reduced expression despite promoter hypomethylation, suggesting that additional regulatory mechanisms are involved. In particular, our study did not examine changes in histone or RNA methylation. For example, *CDKN1A* expression can be regulated by multiple chromatin modifications, including histone acetylation and methylation, which may contribute to its transcriptional regulation alongside DNA methylation (54). DZNep has previously been shown to affect EZH2/PRC2 function and reduce the levels of the repressive histone modification H3K27me3 (31). More broadly, DZNep treatment can alter histone methylation and has been associated with the reactivation of silenced genes (55). Previous studies have also demonstrated relationships between activating H3K4me3, repressive H3K27me3, chromatin accessibility and DNA methylation at TSSs (56). These findings suggest that changes in histone modifications may contribute to the transcriptional responses observed following AHCY inhibition, particularly at genes showing discordant changes in promoter methylation and expression.

Future studies could combine profiling of histone modifications with ATAC-seq to investigate how AHCY inhibition affects chromatin regulation and accessibility. These approaches would be particularly informative at genes showing discordant changes in promoter methylation and expression. In addition, examining different durations of DZNep exposure could help distinguish early responses to AHCY inhibition from longer-term cellular adaptations. Functional studies of candidate regulators such as CDKN1A would also help clarify the mechanisms underlying the reduced proliferation observed following AHCY inhibition.

In summary, our study integrates metabolomic profiling, nanopore-based DNA methylation analysis and RNA-seq to characterize the consequences of AHCY inhibition in colorectal cancer cells. We show that AHCY inhibition induces widespread DNA hypomethylation, alters nucleotide metabolism and transcriptional programs, and reduces cell proliferation. Importantly, promoter hypomethylation was preferentially associated with gene upregulation, including at genes involved in cell cycle regulation such as CDKN1A. Together, these findings highlight the connection between methionine cycle metabolism, DNA methylation and transcriptional regulation, and suggest that altered promoter methylation may contribute to the cellular responses to AHCY inhibition.

## Materials and methods

### Cell lines and cell culturing

Colon cancer cell line GP5d was obtained from Sigma-Aldrich (CAT#). High glucose (4,5 g/ml) DMEM was used for cell culturing. The media was supplemented with 10% fetal bovine serum, 1% penicillin-streptomycin and 2 nM L-glutamine.

### RNA-seq and RT-qPCR

GP5d cells were treated with 1 µM DZNep or vehicle (RNase-free water) for 72 hours, after which the medium was replaced with fresh medium containing DZNep or vehicle. Cells were collected 6 hours later for RNA extraction. Three replicate samples were prepared per condition, and RNA was isolated from each triplicate separately. RNA was isolated for RNA-seq using RNeasy Plus Mini kit (50) (Qiagen) and RNA-libraries were generated using KAPA stranded mRNA-seq kit for Illumina (Roche). RNA was isolated from GP5d cell line both from the control and DZNep treated cells and RNA quality was checked using RNA ScreenTape for Tapestation (Agilent). A total of 500 ng of RNA was used for each library. Libraries were paired-end sequenced using illumina platforms.

RT-qPCR was performed to first test the expression of individual genes and later on as quality control for RNA-seq libraries. For RNA-seq libraries, RT-qPCR was done according to KAPA Library Quantification kit, Illumina Platforms (Roche). qPCR primers for RNA amplification were designed with Primers3 (version 4.1.0; https://primer3.ut.ee/) and tested with UCSC In-Silico PCR (https://genome.ucsc.edu/cgi-bin/hgPcr). RNA-libraries’ qualities were also checked by using D5000 ScreenTape assay for Tapestation (Agilent).

### gDNA isolation and NaNoMe-seq

gDNA of GP5d was first acquired by following QuickExtract protocol (Lucigen) and then isolated with phenol-chloroform-isoamyl alcohol extraction method (PCIAA). The length of gDNA was then measured with D5000 ScreenTape assay for Tapestation (Agilent). Genomic DNA was size-selected to approximately 10 kb fragments using a g-TUBE (520079, Covaris). Sequencing libraries were prepared from 400 ng of genomic DNA using the Native Barcoding Kit (SQK-NBD114.24, Oxford Nanopore Technologies) according to the manufacturer’s instructions. Following adapter ligation, 35 fmol (280 ng) of the multiplexed genomic DNA library was loaded onto the flow cell for sequencing.

NaNoMe-seq was performed on DMSO control and AHCY-inhibited cells as previously described (57). Briefly, nuclei were isolated and treated with the GpC methyltransferase M.CviPI (New England Biolabs, M0227) to methylate accessible chromatin regions. Following GpC methylation, genomic DNA was extracted from nuclei using a phenol-chloroform extraction protocol. Sequencing libraries were prepared from 400 ng of genomic DNA using the Native Barcoding Kit (SQK-NBD114.24, Oxford Nanopore Technologies) according to the manufacturer’s instructions. After adapter ligation, 300 ng of the multiplexed genomic DNA library was loaded onto an Oxford Nanopore flow cell for sequencing.

### Metabolite profiling

GP5d was seeded on day 1 to all wells in a 6-well plate. 3 of the 6 wells in the 6-well plate were then treated with DZNep and the rest were given RNAase free water. The cells were then incubated for 72 hours. On day 4 the media was changed once and DZNep and water were added the same way as before. After 6 hours, the media was aspirated, the cells were washed with PBS and extracted by scraping the cells to ice-cold extraction buffer, and then the cells were sent to Helsinki Metabolomics unit for metabolite profiling. There the samples were sonicated for 60 seconds and vortexed for 120 seconds, both done in three cycles, and then they were centrifuged at 14 000 rpm for 10 minutes at +4°C and evaporated dry under nitrogen stream in evaporation tubes. Then the samples were reconstituted in 40 µl extraction solvent and transferred to LC-MS vials. The samples were then analyzed with LC-MS.

### Proliferation experiment

Cells were seeded on day 0 to 96-well plate. The cells were seeded in 12 wells per condition. The number of cells in each well was 6000 and DZNep and control were the conditions. On day 0, the cells were seeded and they were allowed to attach to the plate for the first 5 hours. After that, DZNep was added to the concentration of 1 µM for DZNep condition. The first 3 wells per condition and 3 empty wells as fluorescence control were tested with fluorescence measurements according to the CyQUANT NF cell proliferation assay kit protocol (Invitrogen) after the cells attached to the plate. The media was changed, new DZNep was added and another 3 wells per condition and 2 empty wells as fluorescence control were tested every second day till day 6.

### Data analyses

FastQC v.0.12.1 (58) was used for quality control. RNA-seq data was first trimmed with fastp v.1.0.1 (59), then the reads were aligned with STAR v.2.7.10b, sorted with Samtools v.1.18 (60) and counted with HTSeq v.2.0.9 (61), and then the analyses were done using DESeq2 in R Statistical Software v.4.4.0, and GSEA (62–64). A read count cutoff of 50 to infinity per gene was used before using DESeq2 to filter out genes with too low expression. For GSEA, Hallmarks gene sets and C2 Reactome subset of CP were primarily used when performing the gene set enrichment analyses. Other gene sets included were from C2 WikiPathways, C2 KEGG Legacy and C5 GO BP. The minimum genes per gene set was set to 10 genes. Other advanced settings for GSEA analysis were not modified.

The NaNOMe data for control and AHCY inhibition data was aligned with Dorado aligner v.0.7.1+80da5f5 (https://github.com/nanoporetech/dorado) using default parameters. The output .bam files were concatenated into one and CpG methylation was processed with modkit v.0.4.4 (https://github.com/nanoporetech/modkit) with parameters --ignore h --motif TCG 1 --motif CCG 1 -- motif ACG 1 to avoid calling methylation on ambiguous GCG trinucleotides in NaNOMe-seq. The modkit CpG bedMethyl files were imported in R v.4.5.2 with a custom script to bsseq objects using bsseq v.1.40.0 (Hansen et al. 2012)(65). The bsseq objects were smoothed with bsseq BSmooth using default parameters. Differentially methylated regions (DMRs) for the BSmooth objects were called using the findDMRs script (https://github.com/timplab/nanopore-methylation-utilities) from Lee et al. (2020) for the AHCY inhibition methylation using the vehicle-treated sample as a control using default parameters (66).

The primary Nanopore sequencing data was viewed using IGV (67) and analyzed further by using R with the help of the description of the modkit used in handling the raw methylation data (68). To detect statistically significant methylation changes globally, we defined significant DMRs as having at least 4 CpGs, CpG methylations being in the center 90% quantile range, an FDR value of less than or equal to 0.05 for methylations, being at least 100 base pairs long and at least 300 base pairs away from other DMRs, and then counted them and grouped them into hypomethylated and hypermethylated DMRs according to direction of change. DMRs were then categorised to promoters, introns, exons and intergenic regions with GRCh38 annotations (GENCODE GRCh38 gene annotations version 44 (Ensembl 110)), and to CpG islands, CpG shores, CpG shelves and open sea with UCSC hg38 annotations from TxDb.Hsapiens.UCSC.hg38.knownGene v.3.20.0 annotation package (69) by using annotatr package v.1.32.0 in R (70). Similarly, to group promoters into significant hypo- and hypermethylations and to compare the promoter methylation change to RNA-seq data, we calculated the average CpG methylation and average methylation change value for each promoter. This was also done to all gene bodies. Average methylation change was considered significant hypomethylation if the average CpG methylation level change was less than −10% and Fisher’s test adjusted p-value was less than 0.05, and similarly the change was considered significant hypermethylation if the change was more than +10% and adjusted p-value less than 0.05. For all analyses with nanopore sequencing data, promoters were defined as areas starting 2500 base pairs upstream from TSS and ending 500 base pairs downstream from TSS. Gene bodies were defined as all the exons and introns and TSS of the genes. The observed vs expected calculation and analysis was done using Monte Carlo simulation as described in Magi et al. (Communications biology, 2023), using 1 000 iterations (71).

To determine whether hypo- or hypermethylated DMRs overlapped PMDs, we used MethylSeekR v.1.50.0 in R (72) with a CpG coverage threshold of ≥5. Bsseq package v.1.46.0 (65) was used to load the bedmethyl output to R. Single nucleotide polymorphisms (SNPs) were called from GP5d control condition nanopore sequencing data with Clair3 v.2.0.3 using default parameters (73) and CpGs overlapping SNPs were removed. To assess the statistical significance of the enrichment of hypo- and hypermethylated DMRs in PMDs, one-sided Fisher’s exact test was used and p-values were adjusted with Benjamini-Hochberg correction.

Metabolite profiling data was analysed using Metaboanalyst and R (74). Before using metaboanalyst, metabolites with pooled quality control RSD over 30%, average blank response (% of pooled quality control) over 20%, metabolites not found or with poor chromatography were filtered out. Then the data was normalized according to the normalizing coefficients provided by Helsinki metabolomics unit. In Metaboanalyst, metabolites with individual missing values were assigned one fifth of the lowest observed value within the same condition, data was transformed to log_10_ and scaled with pareto scaling.

Pheatmap v.1.0.13 (75) was used for heatmaps and ggplot2 v.4.0.0 (76) was used for other plots. Metaboanalyst was used for making the metabolite volcano plot.

## Acknowledgements

We thank HiLIFE research infrastructure for FIMM NGS Genomics laboratory at the University of Helsinki for sequencing services, Helsinki Metabolomics Center for metabolic profiling, and the Center for Scientific Computing (CSC), Finland, for the computational infrastructure. P.P. was supported by Research Council of Finland (356021). B.S. was supported by Research Council of Norway (187615), University of Oslo through the Norwegian Centre for Molecular Biosciences and Medicine (NCMBM) (to Sahu group), Norwegian Cancer Society (274630), South Eastern Norway Health Authority (HSØ) (2025083), Research Council of Finland (320114), Sigrid Jusélius Foundation, Jane and Aatos Erkko Foundation, and iCAN Digital Precision Cancer Medicine Flagship (320185). K.K was supported by the iCANDOC doctoral program. S.D. was supported by the iCANPOD postdoctoral program through the iCANDOC doctoral pilot in precision cancer medicine. We thank Veera Erkkilä and other members of Pihlajamaa and Sahu groups for technical assistance and scientific discussion.

## Author contributions

P.P. conceived and supervised the study. I.T. performed RNA-seq, metabolic profiling and proliferation experiments, and analysis for RNA-seq, metabolomics and downstream analysis for nanopore data. D.P. performed nanopore sequencing experiments and K.K. processed the nanopore data and performed DMR analysis. S.D. supported experimental work and data analysis. B.S. supported the nanopore sequencing and contributed to data interpretation. P.P. and I.T. wrote the manuscript with contributions from all authors.

## Declaration of interests

Authors have no potential conflicts of interest.

## Notes

### Competing Interest Statement

The authors have declared no competing interest.

## References

1. Vizán P, Di Croce L, Aranda S. Functional and Pathological Roles of AHCY. Front Cell Dev Biol. 2021 Mar 31;9. doi:10.3389/fcell.2021.654344

2. Pavičić I, Rokić F, Vugrek O. Effects of S-Adenosylhomocysteine Hydrolase Downregulation on Wnt Signaling Pathway in SW480 Cells. Int J Mol Sci. 2023 Jan;24(22):16102. doi:10.3390/ijms242216102

3. Turner MA, Yang X, Yin D, Kuczera K, Borchardt RT, Howell PL. Structure and function of S-adenosylhomocysteine hydrolase. Cell Biochem Biophys. 2000 Oct 1;33(2):101–25. doi:10.1385/CBB:33:2:101

4. Stanhope SC, Weake VM. AHCY: A metabolic gatekeeper at the interface of methylation, redox balance, and cellular stress response. J Biol Chem. 2026 Feb 2;302(3):111220. doi:10.1016/j.jbc.2026.111220 PubMed PMID: 41638428; PubMed Central PMCID: PMC12955632.

5. Niu S, Zheng X, Yao Y, Dong Y, Hu Y, Xiao Z, et al. The Role of AHCY Expression in Bladder Urothelial Carcinoma: A Bioinformatics and Experimental Analysis. Cancer Manag Res. 2025 Mar 21;17:661–73. doi:10.2147/CMAR.S491044 PubMed PMID: 40144861; PubMed Central PMCID: PMC11937648.

6. Vande Voorde J, Steven RT, Najumudeen AK, Ford CA, Dexter A, Gonzalez-Fernandez A, et al. Metabolic profiling stratifies colorectal cancer and reveals adenosylhomocysteinase as a therapeutic target. Nat Metab. 2023 Aug;5(8):1303–18. doi:10.1038/s42255-023-00857-0

7. Uchiyama N, Dougan DR, Lawson JD, Kimura H, Matsumoto S ichi, Tanaka Y, et al. Identification of AHCY inhibitors using novel high-throughput mass spectrometry. Biochem Biophys Res Commun. 2017 Sep 9;491(1):1–7. doi:10.1016/j.bbrc.2017.05.107

8. Belužić L, Grbeša I, Belužić R, Park JH, Kong HK, Kopjar N, et al. Knock-down of AHCY and depletion of adenosine induces DNA damage and cell cycle arrest. Sci Rep. 2018 Sep 18;8:14012. doi:10.1038/s41598-018-32356-8 PubMed PMID: 30228286; PubMed Central PMCID: PMC6143609.

9. Xie Z, Bi C, Cheong LL, Liu SC, Huang G, Zhou J, et al. Determinants of Sensitivity to DZNep Induced Apoptosis in Multiple Myeloma Cells. PLoS ONE. 2011 Jun 24;6(6):e21583. doi:10.1371/journal.pone.0021583 PubMed PMID: 21720561; PubMed Central PMCID: PMC3123372.

10. Hayden A, Johnson PWM, Packham G, Crabb SJ. S-adenosylhomocysteine hydrolase inhibition by 3-deazaneplanocin A analogues induces anti-cancer effects in breast cancer cell lines and synergy with both histone deacetylase and HER2 inhibition. Breast Cancer Res Treat. 2011 May 1;127(1):109–19. doi:10.1007/s10549-010-0982-0

11. Jiang Q, Lan S, Tan F, Liang Y, Guo Z, Hou Y, et al. Adenosylhomocysteinase plays multiple roles in maintaining the identity and pluripotency of mouse embryonic stem cells†. Biol Reprod. 2024 Mar 1;110(3):450–64. doi:10.1093/biolre/ioad165

12. Bian Y, Li W, Kremer DM, Sajjakulnukit P, Li S, Crespo J, et al. Cancer SLC43A2 alters T cell methionine metabolism and histone methylation. Nature. 2020 Sep;585(7824):277–82. doi:10.1038/s41586-020-2682-1

13. Xu Z, Chen J. A moonlighting function: the methionine cycle rewires RNA methylation. Cell Res. 2026 Apr;36(4):241–2. doi:10.1038/s41422-026-01231-x

14. He PC, He C. m6A RNA methylation: from mechanisms to therapeutic potential. EMBO J. 2021 Feb 1;40(3):e105977. doi:10.15252/embj.2020105977 PubMed PMID: 33470439; PubMed Central PMCID: PMC7849164.

15. Liao K, Cao F, Wei C, Qian ZY, Hu HR, Pan WF, et al. The AHCY–adenosine complex rewires mRNA methylation to enhance fatty acid biosynthesis and tumorigenesis. Cell Res. 2026 Feb;36(2):152–72. doi:10.1038/s41422-025-01213-5

16. Jang HS, Shin WJ, Lee JE, Do JT. CpG and Non-CpG Methylation in Epigenetic Gene Regulation and Brain Function. Genes. 2017 Jun;8(6):148. doi:10.3390/genes8060148

17. Stefansson OA, Sigurpalsdottir BD, Rognvaldsson S, Halldorsson GH, Juliusson K, Sveinbjornsson G, et al. The correlation between CpG methylation and gene expression is driven by sequence variants. Nat Genet. 2024 Aug;56(8):1624–31. doi:10.1038/s41588-024-01851-2

18. Patel D, Tiusanen V, Karttunen K, Pihlajamaa P, Sahu B. Cancer cell type-specific derepression of transposable elements by inhibition of chromatin modifier enzymes. Commun Biol. 2025 Jul 3;8(1):992. doi:10.1038/s42003-025-08413-0

19. Wang P, Chen LL, Xiong Y, Ye D. Metabolite regulation of epigenetics in cancer. Cell Rep. 2024 Oct 22;43(10). doi:10.1016/j.celrep.2024.114815 PubMed PMID: 39368084.

20. Blindu AS, Berardelli S, De Paoli F, Manai F, Tricarico R, Zucca S, et al. Exploring the Impact of DNA Methylation on Gene Expression in CRC: A Computational Approach for Identifying Epigenetically Regulated Genes in Multi-Omic Datasets. Cancers. 2026 Jan 9;18(2):211. doi:10.3390/cancers18020211 PubMed PMID: 41595132; PubMed Central PMCID: PMC12839322.

21. Joe S, Kim J, Lee JY, Jeon J, Byeon I, Han SW, et al. Epigenetic insights into colorectal cancer: comprehensive genome-wide DNA methylation profiling of 294 patients in Korea. BMB Rep. 2023 Oct 31;56(10):563–8. doi:10.5483/BMBRep.2023-0096 PubMed PMID: 37574809; PubMed Central PMCID: PMC10618077.

22. Desterke C, Jaulin F, Dornier E. Hypermethylated Colorectal Cancer Tumours Present a Myc-Driven Hypermetabolism with a One-Carbon Signature Associated with Worsen Prognosis. Biomedicines. 2024 Mar;12(3):590. doi:10.3390/biomedicines12030590

23. Sng CCT, Starling N, Graham TA, Sottoriva A. The genomic and epigenomic evolution of colorectal cancer. Nat Rev Gastroenterol Hepatol. 2026 Nov 2;1–19. doi:10.1038/s41575-026-01265-x

24. WHO. Colorectal cancer [Internet]. [cited 2025 Dec 16]. Available from: https://www.who.int/news-room/fact-sheets/detail/colorectal-cancer

25. Heidelberger C, Chaudhuri NK, Danneberg P, Mooren D, Griesbach L, Duschinsky R, et al. Fluorinated Pyrimidines, A New Class of Tumour-Inhibitory Compounds. Nature. 1957 Mar;179(4561):663–6. doi:10.1038/179663a0

26. Dickinson ME, Flenniken AM, Ji X, Teboul L, Wong MD, White JK, et al. High-throughput discovery of novel developmental phenotypes. Nature. 2016 Sep;537(7621):508–14. doi:10.1038/nature19356

27. Miller MW, Duhl DM, Winkes BM, Arredondo-Vega F, Saxon PJ, Wolff GL, et al. The mouse lethal nonagouti (a(x)) mutation deletes the S-adenosylhomocysteine hydrolase (Ahcy) gene. EMBOJ. 1994 Apr 1;13(8):1806–16. doi:10.1002/j.1460-2075.1994.tb06449.x

28. Aranda S, Alcaine-Colet A, Blanco E, Borràs E, Caillot C, Sabidó E, et al. Chromatin capture links the metabolic enzyme AHCY to stem cell proliferation. Sci Adv. 2019 Mar 6;5(3):eaav2448. doi:10.1126/sciadv.aav2448

29. DepMap, Broad. DepMap Public 26Q3. Dataset. 2026. Available from: https://depmap.org/portal

30. Chera A, Stancu-Cretu M, Zabet NR, Bucur O. Shedding light on DNA methylation and its clinical implications: the impact of long-read-based nanopore technology. Epigenetics Chromatin. 2024 Dec 30;17(1):39. doi:10.1186/s13072-024-00558-2

31. Marquez VE. 3-Deazaneplanocin A (DZNep): A Drug That Deserves a Second Look. J Med Chem. 2024 Oct 11;67(20):17964–79. doi:10.1021/acs.jmedchem.4c01566

32. Arafeh R, Shibue T, Dempster JM, Hahn WC, Vazquez F. The present and future of the Cancer Dependency Map. Nat Rev Cancer. 2025 Jan;25(1):59–73. doi:10.1038/s41568-024-00763-x PubMed PMID: 39468210.

33. Antonioli L, Valdiserra G, Pacher P, Robson SC, Haskó G. Metabolic reprogramming and intracellular ATP homeostasis in immunity. Pharmacol Ther. 2026 May 1;281:109011. doi:10.1016/j.pharmthera.2026.109011

34. Tran DH, Kim D, Kesavan R, Brown H, Dey T, Soflaee MH, et al. De novo and Salvage Purine Synthesis Pathways Across Tissues and Tumors. Cell. 2024 Jul 11;187(14):3602–3618.e20. doi:10.1016/j.cell.2024.05.011 PubMed PMID: 38823389; PubMed Central PMCID: PMC11246224.

35. Camici M, Garcia-Gil M, Tozzi MG. The Inside Story of Adenosine. Int J Mol Sci. 2018 Mar;19(3):784. doi:10.3390/ijms19030784

36. Mathews CK. Deoxyribonucleotides as genetic and metabolic regulators. FASEB J. 2014 Sep;28(9):3832–40. doi:10.1096/fj.14-251249 PubMed PMID: 24928192; PubMed Central PMCID: PMC5395721.

37. MacKenzie M, Argyropoulos C. An Introduction to Nanopore Sequencing: Past, Present, and Future Considerations. Micromachines. 2023 Feb 16;14(2):459. doi:10.3390/mi14020459 PubMed PMID: 36838159; PubMed Central PMCID: PMC9966803.

38. Zhou W, Dinh HQ, Ramjan Z, Weisenberger DJ, Nicolet CM, Shen H, et al. DNA methylation loss in late-replicating domains is linked to mitotic cell division. Nat Genet. 2018 Apr;50(4):591–602. doi:10.1038/s41588-018-0073-4 PubMed PMID: 29610480; PubMed Central PMCID: PMC5893360.

39. Kafetzopoulos I, Taglini F, Pasquier M, Davidson-Smith H, Rodger CJ, Gimeno LP, et al. DNMT1 loss leads to hypermethylation of a subset of late replicating domains by DNMT3A. PLOS Genet. 2026 Apr 2;22(4):e1012098. doi:10.1371/journal.pgen.1012098

40. Wieczorek A, Balwierz W. The Role of Id2 Protein in Neuroblatoma in Children. Pathol Oncol Res. 2015;21(4):999–1004. doi:10.1007/s12253-015-9908-9 PubMed PMID: 25771836; PubMed Central PMCID: PMC4550645.

41. Deng J, Pan T, Liu Z, McCarthy C, Vicencio JM, Cao L, et al. The role of TXNIP in cancer: a fine balance between redox, metabolic, and immunological tumor control. Br J Cancer. 2023 Dec;129(12):1877–92. doi:10.1038/s41416-023-02442-4

42. Heinrichs M, Finke AF, Aibara S, Krempler A, Boshnakovska A, Rehling P, et al. Coupling of ribosome biogenesis and translation initiation in human mitochondria. Nat Commun. 2025 Apr 17;16(1):3641. doi:10.1038/s41467-025-58827-x

43. Nolfi-Donegan D, Braganza A, Shiva S. Mitochondrial electron transport chain: Oxidative phosphorylation, oxidant production, and methods of measurement. Redox Biol. 2020 Aug 6;37:101674. doi:10.1016/j.redox.2020.101674 PubMed PMID: 32811789; PubMed Central PMCID: PMC7767752.

44. Kotrys AV, Szczesny RJ. Mitochondrial Gene Expression and Beyond—Novel Aspects of Cellular Physiology. Cells. 2020 Jan;9(1):17. doi:10.3390/cells9010017

45. Herzig S, Shaw RJ. AMPK: guardian of metabolism and mitochondrial homeostasis. Nat Rev Mol Cell Biol. 2018 Feb;19(2):121–35. doi:10.1038/nrm.2017.95

46. Marin TL, Gongol B, Zhang F, Martin M, Johnson DA, Xiao H, et al. AMPK promotes mitochondrial biogenesis and function by phosphorylating the epigenetic factors DNMT1, RBBP7, and HAT1. Sci Signal. 2017 Jan 31;10(464):eaaf7478. doi:10.1126/scisignal.aaf7478 PubMed PMID: 28143904; PubMed Central PMCID: PMC5830108.

47. Saldivar JC, Cortez D, Cimprich KA. The essential kinase ATR: ensuring faithful duplication of a challenging genome. Nat Rev Mol Cell Biol. 2017 Oct;18(10):622–36. doi:10.1038/nrm.2017.67

48. Mustafa AHM, Krämer OH. Novel insight into mechanisms for ATR activation by chromatin structures. Arch Toxicol. 2021;95(10):3433–4. doi:10.1007/s00204-021-03133-w PubMed PMID: 34382094; PubMed Central PMCID: PMC8448697.

49. Haakonsen DL, Heider M, Ingersoll AJ, Vodehnal K, Witus SR, Uenaka T, et al. Stress response silencing by an E3 ligase mutated in neurodegeneration. Nature. 2024 Feb;626(8000):874–80. doi:10.1038/s41586-023-06985-7

50. Lin K, Wei L, Wang R, Li L, Song S, Wang F, et al. Disrupted methionine cycle triggers muscle atrophy in cancer cachexia through epigenetic regulation of REDD1. Cell Metab. 2025 Feb 4;37(2):460–476.e8. doi:10.1016/j.cmet.2024.10.017 PubMed PMID: 39729999.

51. Fiskus W, Wang Y, Sreekumar A, Buckley KM, Shi H, Jillella A, et al. Combined epigenetic therapy with the histone methyltransferase EZH2 inhibitor 3-deazaneplanocin A and the histone deacetylase inhibitor panobinostat against human AML cells. Blood. 2009 Sep 24;114(13):2733–43. doi:10.1182/blood-2009-03-213496 PubMed PMID: 19638619; PubMed Central PMCID: PMC2756128.

52. Chatterjee B, Ghosh K, Kanade SR. Curcumin-mediated demethylation of the proximal promoter CpG island enhances the KLF4 recruitment that leads to increased expression of p21Cip1 in vitro. J Cell Biochem. 2019;120(1):809–20. doi:10.1002/jcb.27442

53. Wu G, Wang N, Luo Y, Zhang Y, Wang P, Zhu Z, et al. Metabolic perturbation of epigenome by inhibiting S-adenosylhomocysteine hydrolase elicits senescence through DNA damage response in hepatoma cells. Tumor Biol. 2017 May 1;39(5):1010428317699117. doi:10.1177/1010428317699117

54. Ocker M, Al Bitar S, Monteiro AC, Gali-Muhtasib H, Schneider-Stock R. Epigenetic Regulation of p21cip1/waf1 in Human Cancer. Cancers. 2019 Sep;11(9):1343. doi:10.3390/cancers11091343

55. Miranda TB, Cortez CC, Yoo CB, Liang G, Abe M, Kelly TK, et al. DZNep is a global histone methylation inhibitor that reactivates developmental genes not silenced by DNA methylation. Mol Cancer Ther. 2009 Jun 1;8(6):1579–88. doi:10.1158/1535-7163.MCT-09-0013

56. Lee I, Razaghi R, Gilpatrick T, Molnar M, Gershman A, Sadowski N, et al. Simultaneous profiling of chromatin accessibility and methylation on human cell lines with nanopore sequencing. Nat Methods. 2020 Dec;17(12):1191–9. doi:10.1038/s41592-020-01000-7

57. Sahu B, Pihlajamaa P, Zhang K, Palin K, Ahonen S, Cervera A, et al. Human cell transformation by combined lineage conversion and oncogene expression. Oncogene. 2021 Sep;40(36):5533–47. doi:10.1038/s41388-021-01940-0

58. 58. Babraham Bioinformatics - FastQC A Quality Control tool for High Throughput Sequence Data [Internet]. [cited 2026 Mar 23]. Available from: https://www.bioinformatics.babraham.ac.uk/projects/fastqc/

59. Chen S, Zhou Y, Chen Y, Gu J. fastp: an ultra-fast all-in-one FASTQ preprocessor. Bioinformatics. 2018 Sep 1;34(17):i884–90. doi:10.1093/bioinformatics/bty560

60. Danecek P, Bonfield JK, Liddle J, Marshall J, Ohan V, Pollard MO, et al. Twelve years of SAMtools and BCFtools. GigaScience. 2021 Feb 1;10(2):giab008. doi:10.1093/gigascience/giab008

61. Putri GH, Anders S, Pyl PT, Pimanda JE, Zanini F. Analysing high-throughput sequencing data in Python with HTSeq 2.0. Bioinformatics. 2022 May 13;38(10):2943–5. doi:10.1093/bioinformatics/btac166

62. Mootha VK, Lindgren CM, Eriksson KF, Subramanian A, Sihag S, Lehar J, et al. PGC-1α-responsive genes involved in oxidative phosphorylation are coordinately downregulated in human diabetes. Nat Genet. 2003 Jul;34(3):267–73. doi:10.1038/ng1180

63. Subramanian A, Tamayo P, Mootha VK, Mukherjee S, Ebert BL, Gillette MA, et al. Gene set enrichment analysis: A knowledge-based approach for interpreting genome-wide expression profiles. Proc Natl Acad Sci. 2005 Oct 25;102(43):15545–50. doi:10.1073/pnas.0506580102

64. Love MI, Huber W, Anders S. Moderated estimation of fold change and dispersion for RNA-seq data with DESeq2. Genome Biol. 2014;15(12):550. doi:10.1186/s13059-014-0550-8 PubMed PMID: 25516281; PubMed Central PMCID: PMC4302049.

65. Hansen KD, Langmead B, Irizarry RA. BSmooth: from whole genome bisulfite sequencing reads to differentially methylated regions. Genome Biol. 2012 Oct 3;13(10):R83. doi:10.1186/gb-2012-13-10-r83

66. Lee I, Razaghi R, Gilpatrick T, Molnar M, Gershman A, Sadowski N, et al. Simultaneous profiling of chromatin accessibility and methylation on human cell lines with nanopore sequencing. Nat Methods. 2020 Dec;17(12):1191–9. doi:10.1038/s41592-020-01000-7

67. Robinson JT, Thorvaldsdóttir H, Winckler W, Guttman M, Lander ES, Getz G, et al. Integrative genomics viewer. Nat Biotechnol. 2011 Jan;29(1):24–6. doi:10.1038/nbt.1754

68. nanoporetech/modkit [Rust] [Internet]. Oxford Nanopore Technologies; 2026 [cited 2026 Jul 27]. Available from: https://github.com/nanoporetech/modkit

69. Bioconductor Core Team and Bioconductor Package Maintainer. TxDb.Hsapiens.UCSC.hg38.knownGene: Annotation package for TxDb object(s) [Internet]. 2024. doi:10.18129/B9.bioc.TxDb.Hsapiens.UCSC.hg38.knownGene

70. Cavalcante RG, Sartor MA. annotatr: genomic regions in context. Bioinformatics. 2017;33(15):2381–3. doi:10.18129/B9.bioc.annotatr

71. Magi A, Mattei G, Mingrino A, Caprioli C, Ronchini C, Frigè G, et al. High-resolution Nanopore methylome-maps reveal random hyper-methylation at CpG-poor regions as driver of chemoresistance in leukemias. Commun Biol. 2023 Apr 8;6(1):382. doi:10.1038/s42003-023-04756-8

72. Burger L, Gaidatzis D, Schübeler D, Stadler MB. Identification of active regulatory regions from DNA methylation data. Nucleic Acids Res. 2013 Sep;41(16):e155. doi:10.1093/nar/gkt599 PubMed PMID: 23828043; PubMed Central PMCID: PMC3763559.

73. Zheng Z, Li S, Su J, Leung AWS, Lam TW, Luo R. Symphonizing pileup and full-alignment for deep learning-based long-read variant calling. Nat Comput Sci. 2022 Dec;2(12):797–803. doi:10.1038/s43588-022-00387-x

74. Pang Z, Lu Y, Zhou G, Hui F, Xu L, Viau C, et al. MetaboAnalyst 6.0: towards a unified platform for metabolomics data processing, analysis and interpretation. Nucleic Acids Res. 2024 Jul 5;52(W1):W398–406. doi:10.1093/nar/gkae253

75. Raivo Kolde. pheatmap: Pretty Heatmaps [Internet]. 2025. Available from: https://CRAN.R-project.org/package=pheatmap

76. Hadley Wickham. ggplot2: Elegant Graphics for Data Analysis [Internet]. 2016. Available from: https://ggplot2.tidyverse.org

